# Untargeted proteomics identifies plant substrates of the bacterial-derived ADP-ribosyltransferase AvrRpm1

**DOI:** 10.1101/2023.09.25.558804

**Authors:** Simranjit Kaur, Thomas Colby, Domenika Thieme, Carsten Proksch, Susanne Matschi, Ivan Matić, Lennart Wirthmueller

## Abstract

One class of enzymes that plant pathogens employ to manipulate innate immunity and physiology of the infected cells are host-targeted ADP-ribosyltransferases. The bacterial pathogen *Pseudomonas syringae* uses its type III secretion system to inject several effector proteins with ADP-ribosyltransferase activity into plant cells. One of them, AvrRpm1, ADP-ribosylates the plasma membrane-associated RPM1-INTERACTING PROTEIN 4 (RIN4) in *Glycine max* and *Arabidopsis thaliana* to attenuate targeted secretion of defense-promoting compounds. Substrate identification of host-targeted ADP-ribosyltransferases is complicated by the biochemical lability of the protein modification during plant protein extraction and in several cases required prior knowledge on plant immune signaling pathways that are impaired by the ADP-ribosylating type III effector. Using the AvrRpm1-RIN4 pair as a proof-of-concept, we present an untargeted proteomics workflow for enrichment and detection of ADP-ribosylated proteins and peptides from plant cell extracts that in several cases provides site-resolution for the modification.

## Introduction

The ability to limit activation of plant innate immunity is a pre-requisite for successful infection by microbial pathogens (Wang et al., 2022). The Gram-negative bacterial pathogen *Pseudomonas syringae* pv. *tomato* (*Pst*) injects 29 type III effectors (T3Es) into the host cell cytoplasm. Collectively, these effectors manipulate host physiology and suppress plant immune responses triggered by recognition of microbe-associated molecular patterns (MAMPs) (Xin et al., 2016; Cunnac et al., 2011). MAMP recognition by cell surface localized pattern recognition receptors activates pattern-triggered immunity (PTI) that is particularly effective against attempted invasion by non-adapted pathogens. In contrast, well-adapted pathogens effectively subvert PTI by translocating effectors into plant cells (Wang et al., 2022). Immunity against such adapted pathogens in plants is conferred by a second class of intracellular immune receptors, nucleotide-binding domain leucine-rich repeat receptors (NLRs), in a cultivar-specific manner (Saur et al., 2021). This form of NLR-mediated resistance to specific pathogen isolates is known as effector-triggered immunity (ETI) (Saur et al., 2021). NLRs detect pathogen effectors either directly or they form protein complexes with virulence targets of effectors and sense effector-mediated manipulation of the ‘guarded’ plant proteins (Wang et al., 2019; Martin et al., 2020). Therefore, the outcome of an infection attempt by an adapted pathogen depends on both, the set of effectors delivered by the pathogen and the NLR repertoire of the plant. In the absence of NLR-mediated recognition effectors promote pathogen virulence (Wang et al., 2022). In contrast, in interactions where at least one effector is recognized by an NLR, the plant rapidly activates ETI that limits further pathogen spread and often involves localized programmed cell death at the site of infection (Saur et al., 2021).

Many bacterial T3Es are host-targeted enzymes that manipulate plant proteins by post-translational modifications to perturb activation of plant immunity (Wang et al., 2022). Among the T3Es that *Pst* strain DC3000 injects into plant cells are several ADP-ribosyltransferases that modify host proteins by transfer of the ADP-ribose (ADPr) moiety from nicotinamide adenine dinucleotide (NAD^+^) onto amino acid side chains (Fu et al., 2007; Wang et al., 2010; Aung et al., 2020; Suskiewicz et al., 2023). The T3E HopF2 ADP-ribosylates Arabidopsis MITOGEN-ACTIVATED PROTEIN KINASE KINASE 5 (MKK5), thereby interfering with signal transduction from activated pattern recognition receptors to the nucleus (Wang et al., 2010). HopF2 can also ADP-ribosylate Arabidopsis RIN4 *in vitro* (Wang et al., 2010). Over-expression of a HopF2 transgene in Arabidopsis prevents cleavage of RIN4 by another T3E, the Cysteine protease AvrRpt2 (Wilton et al., 2010). This indicates that HopF2 ADP-ribosylates residues in the nitrate-induced (NOI) domains of RIN4 that are essential for cleavage by AvrRpt2. RIN4 is targeted by additional T3Es from different *P. syringae* pathovars and it is guarded against effector-induced manipulation by the NLRs RESISTANCE TO P. SYRINGAE PV MACULICOLA 1 (RPM1) and RESISTANT TO P. SYRINGAE 2 (RPS2) in the Col-0 accession (Belkhadir et al., 2004). The T3E AvrRpm1 from *P. syringae* pv. *maculicola* ADP-ribosylates RIN4 proteins from Arabidopsis and soybean at two positions in their N- and C-terminal NOI domains (Redditt et al., 2019). In addition to RIN4, AvrRpm1 also modifies several sequence-related Arabidopsis NOI proteins by ADP-ribosylation (Redditt et al., 2019). Following identification of the ADP-ribosylation sites on soybean RIN4b by proteomics, Redditt et al. (2019) predicted, based on sequence homology, that AvrRpm1 modifies Arabidopsis RIN4 on N11 and D153. RPM1 is activated by modifications in the C-terminal NOI (C-NOI) of RIN4 (Chung et al., 2011). A RIN4 D153A mutant variant that can no longer be ADP-ribosylated in the C-NOI attenuates AvrRpm1-induced activation of RPM1 (Redditt et al., 2019).

Previously, different approaches have been employed to identify substrates of plant-targeted bacterial ADP-ribosyltransferases. Fu et al. (2007) used ADP-ribosylation assays with ^32^P-NAD^+^ in plant cell extracts in combination with anion exchange chromatography and two-dimensional gel electrophoresis to identify the RNA-binding protein GRP7 as a substrate of the T3E HopU1 from *Pst* DC3000. In other cases, prior knowledge about T3E interference with distinct immune signaling pathways, protein interactors and/or subcellular localization of bacterial ADP-ribosyltransferases in plant cells has aided in identification of potential host targets (Wang et al., 2010; Wilton et al., 2010; Zhou et al., 2014; Aung et al., 2020; Wu et al., 2024). For three plant substrates, ADPr modification sites have been determined by targeted *in vitro* ADP-ribosylation assays in combination with site-directed mutagenesis (Fu et al., 2007; Wang et al., 2010; Wu et al., 2024). Detection of ADP-ribosylation sites by proteomics following immunoprecipitation of presumed host target proteins from plant cell extracts has only recently been established and this approach so far relies on prior identification of effector targets and over-expression of epitope-tagged proteins (Redditt et al., 2019; Yoon et al., 2022). Given that putative host-targeted ADP-ribosyltransferases are not limited to *P. syringae* but also present in other bacterial and fungal plant pathogens (Seong and Krasileva, 2021) we sought to establish proteomics methods that can aid in direct detection of ADP-ribosylated proteins from plant protein extracts. Here we present a proteomics workflow that can identify ADP-ribosylated proteins from transgenic lines expressing bacterial ADP-ribosyltransferases by comparative proteomics and provide site resolution for the ADP-ribosylated peptides.

## Results

We used transgenic lines that express AvrRpm1-HA or HopF2-HA under control of a Dexamethasone (Dex)-inducible promoter to assess if ADP-ribosylated endogenous RIN4 protein (24 kDa) can be detected by immunoblot analysis. Samples from two independent AvrRpm1-expressing lines showed two bands between 25 and 35 kDa when probed with an antibody specific to mono-ADP-ribosylation (Bonfiglio et al., 2020). In contrast, the antibody did not detect prominent bands in the two lines expressing HopF2-HA (Fig. 1A). Even upon longer exposure, the band pattern of the HopF2 samples looked similar to those from Col-0 and two lines carrying a transgene encoding the enzymatically inactive HopF2 D175A variant. As shown by the α-HA immunoblot in Fig. 1A, both HopF2-HA protein variants were expressed suggesting that HopF2 substrates might be proteins of low abundance or that they were not efficiently extracted from plant tissue by our method. Next, we used the engineered version of the Af1521 Macro domain (Nowak et al., 2020) to enrich ADP-ribosylated proteins from plant extracts (Fig. 1B). As the Af1521 Macro domain can also bind to the terminal ADPr of poly(ADPr) chains, the signal on the immunoblot probed with α-pan-ADPr binding reagent might represent a mixture of mono- and poly-ADP-ribosylated proteins. When analyzed by LC-MS/MS we identified 24 peptide spectra matches (PSM) of RIN4 in two samples from an AvrRpm1-expressing line but not in Col-0 or HopF2 samples (Table 1, Supplementary Dataset S1). However, this approach did not detect any ADP-ribosylated peptides of RIN4 or other proteins. We found that a depletion of potentially competing nucleotide analogs (Karras et al., 2005) by a Sephadex G-25 column resulted in stronger enrichment by the Af1521 Macro domain (Fig. 1C).

**Figure 1.**
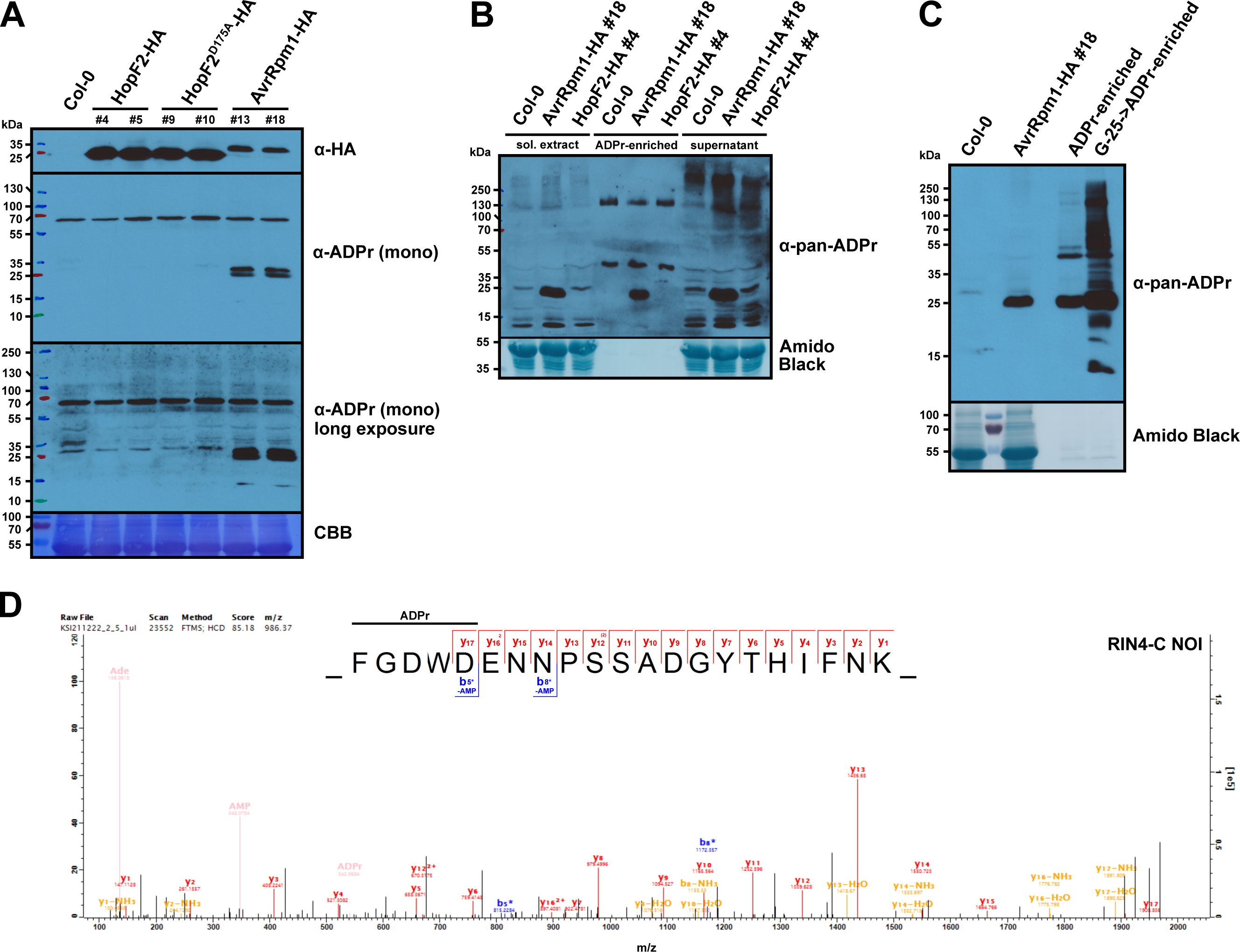
ADP-ribosylated substrates of AvrRpm1 but not HopF2 can be detected by immunoblots, enriched by the engineered Af1521 Macro domain, and identified by untargeted proteomics. **(A)** Detection of AvrRpm1 substrates by an α-ADPr immunoblot. Protein extracts from Dex-treated Col-0 wild type and the indicated transgenic lines were separated by SDS-PAGE, electroblotted onto PVDF membrane and probed with α-HA and α-mono-ADPr antibodies. CBB = Coomassie Brilliant Blue-stained membrane as loading control. **(B)** Enrichment of ADP-ribosylated proteins from the indicated Arabidopsis lines by the engineered Af1521 Macro domain and detection in an α-pan-ADPr immunoblot. Approximately 15% of the ADPr-enriched fraction was loaded onto the gel. ‘sol. extract’ = soluble protein extract. The membrane was stained with Amido Black as loading control. This experiment was repeated once with similar results. **(C)** Depletion of potentially competing nucleotide analogs from plant extracts prior to the Af1521 affinity purification facilitates enrichment of ADP-ribosylated proteins. ‘Col-0’ and ‘AvrRpm1-HA #18’ indicate the respective protein extracts before ADPr-enrichment. ‘ADPr-enriched’ = AvrRpm1 sample, direct pulldown with the Af1521 Macro domain. ‘G-25->ADPr-enriched’ = AvrRpm1 sample, pre-clearing by G-25 sepharose followed by pulldown with the Af1521 Macro domain. ADP-ribosylated proteins were detected as in (B). **(D)** HCD MS^2^ spectrum of the ADP-ribosylated peptide FGDWDENNPSSADGYTHIFNK from the RIN4 C-NOI identified by untargeted proteomics (z = 3, mass = 2954.09 Da). Unmodified fragment ions and fragment ions with a neutral loss of AMP are indicated in the sequence logos. The horizontal line designates the possible sequence window for ADP-ribosylation. The ADPr marker ions are shown in beige. Asterisks indicate b-ions with a neutral loss of AMP (Δ347.06 Da).

**Table 1.** Proteins specifically identified in Af1521 Macro-enriched AvrRpm1-HA samples (≥ 10 PSM) are not or not strongly enriched by the Af1521 G42E mutant variant.

| AGI identifier | description | PSM | PSM | PSM | PSM ratio |
| --- | --- | --- | --- | --- | --- |
|  |  | Col-0 | AvrRpm1 | HopF2 | Af1521 / G42E |
| AT4G11840.1 | PLDGAMMA3 | 0 | 35 | 0 | 28 / 0 |
| AT3G45780.1 | PHOT1 | 0 | 29 | 0 | 27 / 0 |
| AT5G54110.1 | MAMI | 0 | 27 | 0 | 11 / 0 |
| AT3G25070.1 | RIN4 | 0 | 24 | 0 | 27 / 6 |
| AT3G24550.1 | PERK1 | 0 | 18 | 0 | 5 / 0 |
| AT1G70940.1 | PIN3 | 0 | 14 | 0 | 0 / 0 |
| AT5G39590.1 | TLD-domain containing nucleolar protein | 0 | 12 | 0 | 6 / 0 |
| AT1G30470.1 | SIT4 phosphatase-associated family protein | 0 | 11 | 0 | 4 / 0 |
| AT5G53000.1 | TAP46 | 0 | 10 | 0 | 22 / 4 |

To further optimize detection of ADP-ribosylated peptides we focused on substrates of AvrRpm1. Untargeted proteomics of the pre-cleared and ADPr-enriched AvrRpm1 sample detected ADP-ribosylated peptides of the NOI sequences from NOI4 (AT5G55850.3), NOI12 (AT2G04410.1), as well as the N-NOI and C-NOI sequences of RIN4 (Supplementary Dataset S2). Although the marker ions of ADPr fragmentation (Ade^+^ 136.06, AMP^+^ 348.07, ADPr^+^ 542.07 Da) and MaxQuant (Tyanova et al., 2016) Andromeda scores between 64 and 156 clearly identified these peptides as ADP-ribosylated (Fig. 1D), the localization probability for ADPr modification sites was often below 80% and therefore ambiguous. Next, we used a data-dependent ‘triggering’ method that initiates an MS^2^ scan with an elevated threshold for automatic gain control, higher resolution, and longer injection time upon detection of the 136.06 Da Adenine^+^ fragment of ADPr that is typically an intense fragment ion from ADP-ribosylation sites (Rosenthal et al., 2015; Geiszler et al., 2023) (Supplementary Methods). This method resulted in more informative ion series including additional neutral losses of AMP (Δ347.06 Da) (Fig. 2, Supplementary Dataset S3). Consequently, the ADPr localization probabilities improved to >80% and from three LC-MS/MS runs we could identify at least one MS^2^ spectrum for each modified peptide with a localization score of >99.5%. Based on these data and the assumption that ADP-ribosylation does not occur on F, G, or W (Rack et al., 2020), the NOI4 protein was ADP-ribosylated on E29 (Fig. 2A) and for NOI12 we observed ADP-ribosylation on the sequence-equivalent position E13 (Fig. 2B). RIN4 was ADP-ribosylated on N11 (N-NOI, Fig. 2C) and D153 (C-NOI, Fig. 2D). ADP-ribosylation of these residues was further supported by manual inspection of MS^2^ spectra exported from Proteome Discoverer that revealed presence of the respective b3* ions with a neutral loss of AMP (Supplementary Fig. S1). In conclusion, based on the AvrRpm1-RIN4 proof-of-concept experiments, our untargeted method robustly identified RIN4 and RIN4 homologs as *in planta* substrates of AvrRpm1 and allowed mapping the ADP-ribosylation sites with high confidence.

**Figure 2.**
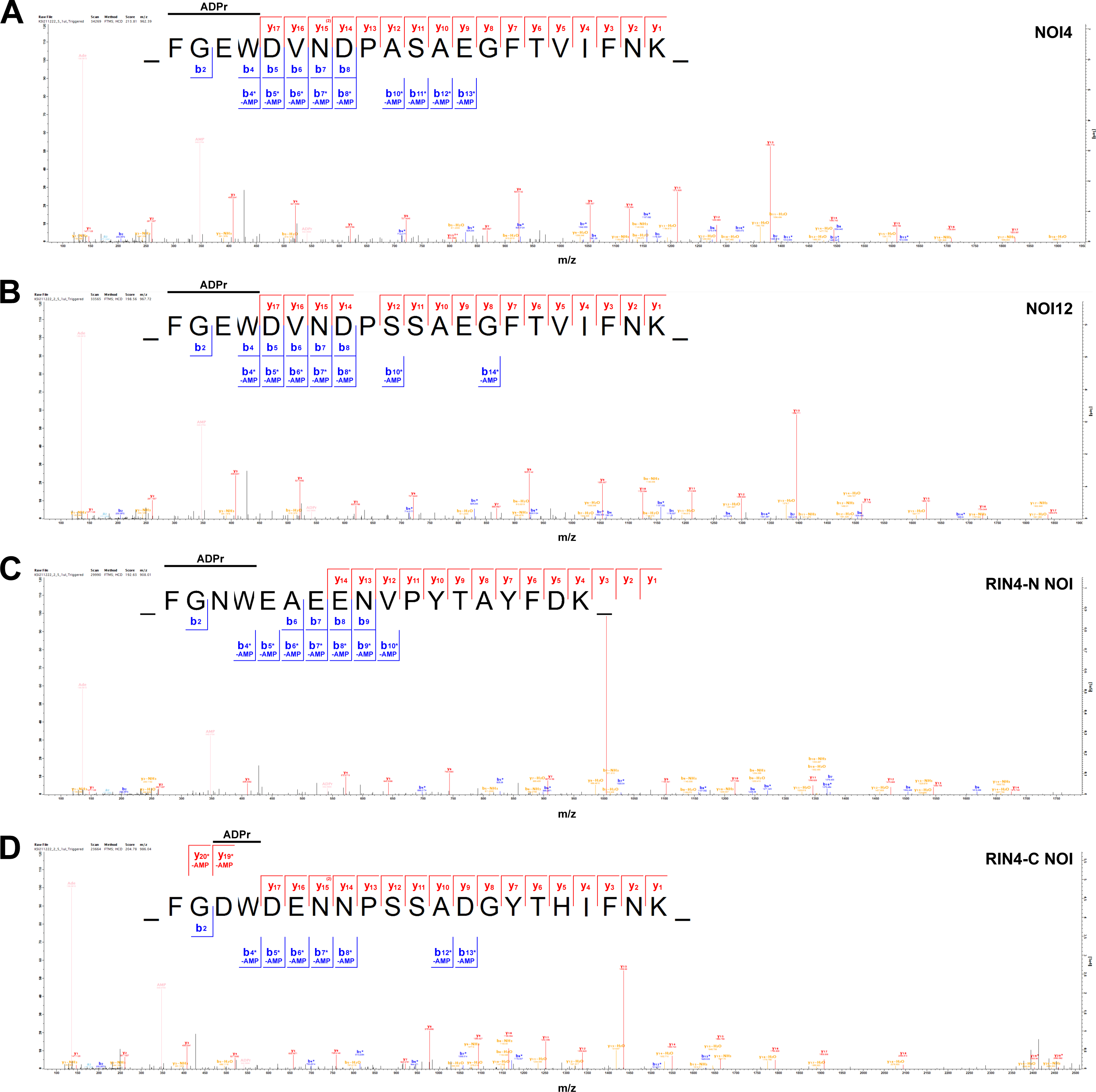
ADPr-modification sites on AvrRpm1 substrates revealed by the ‘triggering’ HCD fragmentation method. Panels A-D show MS^2^ spectra of ADP-ribosylated peptides from NOI4, NOI12, and the RIN4 N-NOI and C-NOI sequences. Unmodified fragment ions and fragment ions with a neutral loss of AMP are indicated in the sequence logos. The horizontal line designates the possible sequence window for ADP-ribosylation. **(A)** NOI4-derived peptide FGEWDVNDPASAEGFTVIFNK (z = 3, mass = 2883.17 Da). **(B)** NOI12-derived peptide FGEWDVNDPSSAEGFTVIFNK (z = 3, mass = 2899.14 Da). **(C)** RIN4 N-NOI-derived peptide FGNWEAEENVPYTAYFDK (z = 3, mass = 2720.01 Da). **(D)** RIN4 C-NOI-derived peptide FGDWDENNPSSADGYTHIFNK (z = 3, mass = 2954.09 Da). The ADPr marker ions are shown in beige. Asterisks indicate ions with a neutral loss of AMP (Δ347.06 Da).

While MaxQuant reports several ADP-ribosylated peptides in addition to those from NOI4, NOI12, and RIN4, their MS^2^ spectra overall had lower Andromeda scores and lacked the ADP-ribosylation marker ions (Supplementary Dataset S3). These spectra were also characterized by a higher signal to noise ratio and in many cases the incomplete b- and y-ion series did not cover the assumed ADP-ribosylation site. Therefore, the data did not provide sufficient evidence for ADP-ribosylation. One exception was a peptide from the MEMBRANE-ASSOCIATED MANNITOL-INDUCED (MAMI, AT5G54110.1) protein (Fig. 3A) that was identified in one of the three LC-MS/MS runs. In this case an almost complete y-ion series with neutral losses of AMP together with the Ade^+^ and AMP^+^ marker ions indicated ADP-ribosylation of peptide VVGEGLVID<u>E</u>WK<u>E</u>R (236-249) with the highest localization probabilities for E245 or E248 (underlined, 33% respectively). Manual inspection of the MS^2^ spectrum revealed a better fit of the recorded spectrum to a peptide with ADPr on E245 (Supplementary Fig. S2). If MAMI is ADP-ribosylated by AvrRpm1, the protein should be enriched compared to controls in our ADPr affinity purification. Indeed, we identified MAMI with 27 PSM in ADPr-enriched samples of Dex-treated AvrRpm1 plants but not in Dex-treated HopF2 or Col-0 controls (Table 1, Supplementary Dataset S1). Over-expression of AvrRpm1 may stabilize or elevate MAMI protein levels, which could result in higher carry-over of MAMI-derived peptides during the affinity enrichment. However, we consider this unlikely as we did not observe PSM for MAMI when we performed the enrichment with the Af1521 G42E variant (Dani et al., 2009) that does not bind ADPr (0 PSM G42E vs. 11 PSM Af1521; Table 1, Supplementary Dataset S4). We obtained further evidence for AvrRpm1-mediated ADP-ribosylation of MAMI from the transient *Nicotiana benthamiana* expression system. When AvrRpm1-Flag was co-expressed with a HA-tagged MAMI we observed an additional slower migrating band on the α-HA immunoblot (Fig. 3B). A band of ∼37 kDa was also visible when the blot was probed with α-pan-ADPr (Fig. 3B), indicating that MAMI can be ADP-ribosylated by AvrRpm1. In LC-MS/MS analysis of samples from *N. benthamiana* leaves co-expressing AvrRpm1-Flag and HA-MAMI we detected the ADP-ribosylated MAMI peptide shown in Fig. 3A only in the pulldown with the functional Af1521 Macro domain but not the G42E control. Expression of HA-MAMI alone did not result in detectable ADP-ribosylation indicating that co-expression of AvrRpm1 was required for ADP-ribosylation of MAMI (Supplementary Dataset S5).

**Figure 3.**
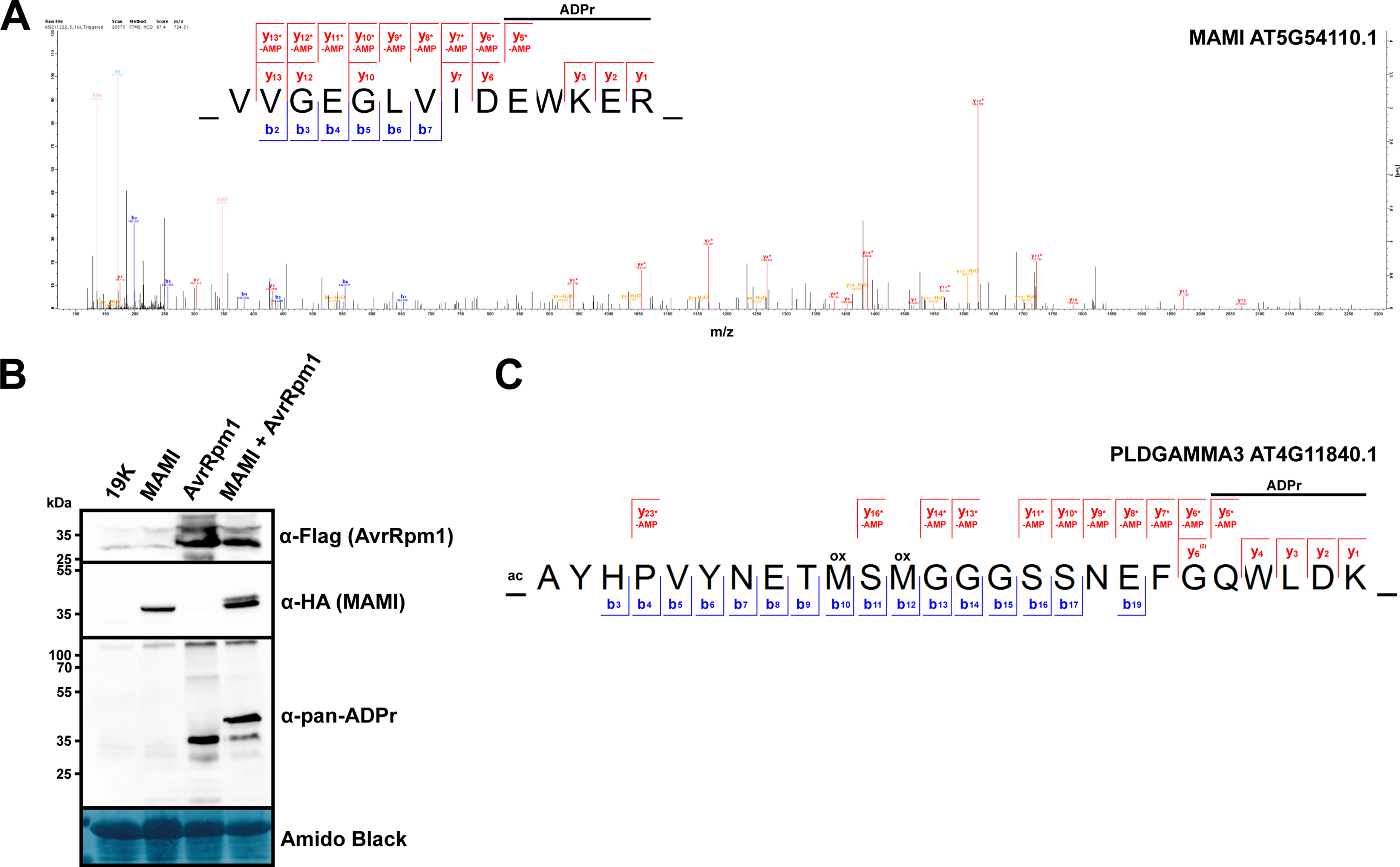
AvrRpm1 ADP-ribosylates Arabidopsis MAMI and PLDGAMMA 3. **(A)** HCD MS^2^ spectrum of the ADP-ribosylated MAMI peptide VVGEGLVIDEWKER (z = 3, mass = 2168.92 Da). The ADPr marker ions are shown in beige. Unmodified y-ions and y-ions with a neutral loss of AMP are indicated above the sequence logo. The horizontal line designates the possible sequence window for ADP-ribosylation. Asterisks indicate ions with a neutral loss of AMP (Δ347.06 Da). **(B)** Detection of ADP-ribosylated StrepII-HA-MAMI transiently co-expressed with AvrRpm1-Flag in *N. benthamiana*. Protein extracts were separated by SDS-PAGE, electroblotted onto PVDF membrane and probed with α-HA, α-pan-ADPr, or α-Flag antibodies. The membrane was stained with Amido Black as loading control. ‘19K’ indicates a sample from leaves expressing only the silencing suppressor 19K. The results are representative of three independent experiments. Images of the complete membranes are shown in Supplementary Figure S3. **(C)** ADP-ribosylated peptide AYHPVYNETMSMGGGSSNEFGQWLDK (z = 3, mass = 3519.31 Da) from PLDGAMMA3-HA-StrepII co-expressed with AvrRpm1-Flag in *N. benthamiana*. Unmodified y-ions and y-ions with a neutral loss of AMP are indicated above the sequence logo. The horizontal line designates the possible sequence window for ADP-ribosylation. Asterisks indicate ions with a neutral loss of AMP (Δ347.06 Da).

Apart from RIN4, NOIs, and MAMI several other proteins were enriched in ADPr pulldowns from AvrRpm1-expressing plants in a G42-dependent manner and were not identified in Col-0 and HopF2 samples. This set includes PHOSPHOLIPASE D GAMMA 3 (PLDGAMMA3, AT4G11840.1) and the blue light photoreceptor PHOTOTROPIN 1 (PHOT1, AT3G45780.1) (Table 1; Supplementary Datasets S1, S4). Therefore, even if no ADP-ribosylated peptides can be identified, comparative pulldowns with the functional vs. non-functional Af1521 Macro domains can provide information on potentially modified proteins that can be further assessed for evidence of ADP-ribosylation by directed methods. We attempted to identify ADP-ribosylated peptides of PHOT1 and PLDGAMMA3 by co-expressing these proteins with AvrRpm1-Flag in *N. benthamiana*, followed by affinity enrichment of ADP-ribosylated proteins and detection by LC-MS/MS. While we did not identify any ADP-ribosylated peptides from PHOT1, profiling of samples co-expressing PLDGAMMA3-HA and AvrRpm1-Flag revealed the N-terminal acetylated peptide shown in Fig. 3C from the phospholipase with Q23 as most likely ADP-ribosylation site (Supplementary Dataset S6, see Supplementary Fig. S4 for an annotated MS^2^ spectrum).

To assess if the ADPr profiling method is applicable to other host-targeted ADP-ribosyltransferases from *Pst*, we extended our analysis to transgenic Arabidopsis lines that conditionally express the T3E HopU1. HopU1 ADP-ribosylates the RNA binding protein GRP7 and two Arginine residues (R47, R49) are essential for HopU1-mediated ADP-ribosylation (Fu et al., 2007). ADP-ribosylation on R49, a residue with an important role in RNA binding (Schöning et al., 2007), has been confirmed by *in vitro* enzyme assays with recombinantly expressed proteins followed by proteomics (Jeong et al., 2011). Dex-induced expression of HopU1-Flag resulted in three prominent bands detected on an α-pan-ADPr immunoblot (Supplementary Fig. S4). The band migrating just above the 15 kDa marker would be consistent with the molecular weight of GRP7 (16 kDa). We applied the ADPr profiling method to plants expressing HopU1-Flag and compared them to Col-0 plants. We identified the ADP-ribosylated GRP7 peptide <u>SR</u>GFGFVTFKDEK (48-60) with a possible modification on S48 or R49 (Supplementary Fig. S4). The modified peptide was specifically detected in the Af1521-enriched samples and not in the G42E controls or Col-0 samples (Supplementary Dataset S7). We conclude that the ADPr profiling method is applicable to different T3Es as long as they induce sufficient levels of plant protein ADP-ribosylation when expressed as a transgene (Supplementary Fig. S4).

## Discussion

In this study, we present a proteomics workflow that, regardless of prior knowledge, would have identified RIN4 and sequence-related NOI proteins as substrates of the host-targeted bacterial ADP-ribosyltransferase AvrRpm1 in Arabidopsis. The method identified the two ADP-ribosylation sites on the endogenous RIN4 protein that were predicted by Redditt et al. (2019) based on LC-MS/MS analysis of the soybean RIN4b protein. We further show that RIN4 homologs NOI4 and NOI12 are ADP-ribosylated at the sequence-equivalent site of the conserved NOI sequence indicating that AvrRpm1 preferentially modifies NOI proteins on Glu, Asp or Asn residues on the 5^th^ position of the PKFG[E/D/N]W[E/D/N] consensus sequence. AvrRpm1 and the sequence-unrelated T3E AvrB indirectly promote RIN4 phosphorylation on T166 by receptor-like cytoplasmic kinases and it is the AvrRpm1-mediated manipulation of the RIN4 C-NOI sequence that is sensed by RPM1(Liu et al., 2011; Chung et al., 2011). Based on single and double mutant analysis of RIN4 D153 and T166, Redditt et al. (2019) hypothesized that ADP-ribosylation on D153 promotes phosphorylation of T166. In contrast, data from Yoon et al. (2022) indicate that AvrRpm1-mediated ADP-ribosylation of a non-phosphorylatable RIN4 T166A mutant protein still triggers RPM1 activation. While we identified several previously reported RIN4 phosphorylation sites (S116, S141, and the peptide SSGANVSGS<u>S</u>R<u>T</u>PTHQSSR with phosphorylation on S186 or T188, Supplementary Dataset S8) (Willems et al., 2019), we did not detect phosphorylation on T166. However, given the relatively low abundance of ADP-ribosylated peptides from the RIN4 C-NOI, we consider this result inconclusive with respect to either of the two hypotheses for RPM1 activation.

In the absence of NLR recognition, i. e. when AvrRpm1-expressing *P. syringae* bacteria are directly infiltrated into leaves of the *rpm1 rps2* double mutant, AvrRpm1 promotes bacterial virulence and this does not require RIN4 (Belkhadir et al., 2004). In this respect, plasma membrane-associated proteins like MAMI, PLDGAMMA3 or PHOT1 that are specifically ADPr-enriched from AvrRpm1-expressing plants might constitute potential virulence targets of AvrRpm1. The PHOT1 orthologs of potato and the liverwort *Marchantia polymorpha* have recently been identified as negative regulators of immunity and the potato late blight pathogen *Phytophthora infestans* employs an effector protein to manipulate immune signaling downstream of Stphot1 (He et al., 2018; Naqvi et al., 2022; Yotsui et al., 2023). These findings may support a role of Arabidopsis PHOT1 as a virulence target of AvrRpm1. However, given that the Dex-induced AvrRpm1 effector is over-expressed in our experimental system, this hypothesis needs to be tested by bacterial growth assays in a *rpm1 rps2 phot1* triple mutant background.

In contrast to AvrRpm1 and HopU1, our approach did not identify substrates of HopF2. Multiple host targets of HopF2 have been proposed including MKK5, RIN4, and BAK1 (Wang et al., 2010; Wilton et al., 2010; Zhou et al., 2014). However, none of these proteins were enriched in ADPr affinity pulldowns from HopF2-expressing plant cell extracts (Supplementary Dataset S1). One possible explanation is that proteins targeted by HopF2 have low abundance, are not effectively extracted by our protein isolation method and/or are rapidly degraded upon modification. However, at least for the proposed HopF2 target RIN4 we can exclude inappropriate extraction conditions or low abundance, as proven by the successful affinity enrichment of ADP-ribosylated RIN4 from transgenic lines expressing AvrRpm1. The Af1521 Macro domain has been used for proteome-wide profiling of ADP-ribosylation sites in mammalian cells and does not show strong preferences in terms of the modified amino acid (Martello et al., 2016; Hendriks et al., 2019). Wang et al. (2010) reported that a constitutively active MKK5 variant is more efficiently ADP-ribosylated by HopF2, suggesting that the kinase might require prior activation for effector-induced modification. At least *in vitro*, we were able to reproduce the HopF2-dependent ADP-ribosylation of MKK5 by using an α-ADPr antibody (Supplementary Fig. S5). We consider the differential efficiency of MKK5 ADP-ribosylation in planta and *in vitro* a possible reason for the lack of ADPr peptide enrichment from plants expressing HopF2.

We envisage that the method presented here will facilitate untargeted proteomics approaches to identify substrates of ADP-ribosyltransferases in plants. LC-MS/MS profiling of proteins that are enriched by the Af1521 Macro domain but not the G42E mutant can provide information on plant proteins that are ADP-ribosylated by a particular effector by comparison to suitable controls. The ‘triggering’ proteomics method has the potential to directly sequence modified ADPr-enriched peptides as demonstrated for NOI4, NOI12, RIN4, MAMI, and GRP7. As shown for MAMI and PLDGAMMA3, the transient *N. benthamiana* expression system can aid in pinpointing the ADP-ribosylated peptide and/or amino acid. Ideally, an efficient and specific enrichment by the Af1521 Macro domain, paired with detection of ADP-ribosylated peptides, will facilitate identification of proteins modified by host-targeted ADP-ribosyltransferases from pathogens.

## Materials and Methods

### Experimental design and statistical rationale

The immunoblots presented in Fig. 1A-C are representative of two sets of samples derived from independent experiments. For Supplementary Dataset S1, samples from two independent experiments were analyzed by LC-MS/MS. Wild-type Col-0 plants served as a control for T3E-dependent ADP-ribosylation. To minimize biological and sample variation during method optimization, Supplementary Datasets S2 and S3 were derived from one biological Dex-AvrRpm1 sample that was subjected to ADPr-enrichment using 1, 2.5, or 5 μg of the Af1521 Macro domain. The derived peptide samples were first analyzed by the standard LC-MS/MS method (Supplementary Dataset S2) and immediately afterwards profiled again using the ‘triggering’ method (Supplementary Dataset S3) to allow for a direct comparison of the two methods. The comparison of proteins enriched by the functional Af1521 Macro domain but not by the G42E variant (Supplementary Dataset S4) was derived from a single Dex-AvrRpm1 sample that was analyzed in two technical replicates per pulldown. Data from Supplementary Datasets S1 and S4 were combined in Table 1 to identify potential AvrRpm1 substrates supported by evidence from both approaches. AvrRpm1-dependent ADP-ribosylation of MAMI (Fig. 3B) was tested in three independent experiments, one of which was profiled by LC-MS/MS (Supplementary Dataset S5) to confirm that the α-pan-ADPr immunoblot signal is indeed due to ADP-ribosylation. The ADP-ribosylated PLDGAMMA3 peptide (Supplementary Dataset S6) was identified from a single experiment. For Supplementary Dataset S7, samples from two independent experiments were analyzed by LC-MS/MS. For proteomics analyses a decoy database search was performed to determine the peptide spectral match (PSM) and peptide identification false discovery rates (FDR). Peptides with a score surpassing the false discovery rate threshold of 0.01 (q-value<0.01) were considered positive identifications. A detailed description of the proteomics methods is provided in Supplementary Methods and Supplementary Dataset S9.

### Plant material, DNA constructs, bacterial strains, and growth conditions

The transgenic lines expressing Dex-inducible AvrRpm1-HA and HopF2-HA have been described (Mackey et al., 2002; Wilton et al., 2010). Transgenic Arabidopsis lines expressing a Dex-inducible HopU1-3xFlag construct were generated by transforming Col-0 plants with pTA7001/des/HopU1-3xFlag. Arabidopsis plants were grown in controlled environment chambers with a 10 h light period and 22/18 °C day/night temperatures. To induce expression of T3Es, 4-week-old plants were sprayed with a solution containing 20 μM Dex and 0.01% Silwet-77 and leaves were harvested 16-18 h later. *N. benthamiana* plants were cultivated under long day (16 h light, 8 h dark) conditions at 22 °C and approximately 30% relative humidity in a greenhouse under supplementary light from Tungsten lamps at approximately 200 μmol s^-1^ m^-^². The G42E mutation in the engineered Af1521 Macro domain was introduced by site-directed mutagenesis in pGEX6P-1 (Nowak et al., 2020). The functional (K35E, Y145R) and non-functional (K35E, G42E, Y145R) variants were cloned into KpnI/HindIII-linearized pOPINF (Berrow et al., 2007) by Gibson cloning to create His6-GST-Macro constructs. *E. coli* expression constructs for HopF2 and MKK5 were created by cloning the coding sequences into KpnI/HindIII-linearized pOPINF (Berrow et al., 2007). The MAMI, PHOT1, and AvrRpm1 coding sequences as well as the genomic sequence of PLDGAMMA3 were cloned into pENTR4 by Gibson cloning. The N-terminally tagged StrepII-3xHA-MAMI construct was created by a Gateway LR reaction with pENS-HS-GW. The C-terminally tagged PHOT1-3xHA-StrepII and PLDGAMMA3-3xHA-StrepII constructs were created by Gateway LR reactions with pXCSG-GW-HS. AvrRpm1 and HopU1 were fused to a 3xFlag tag by a Gateway LR reaction with pTA7001/des/3XFlag (Li et al., 2013). Binary vectors were transformed into *Agrobacterium tumefaciens* strain GV3101 pMP90^RK^ (pENS, pXCSG) or GV3101 pMP90 (pTA7001).

### Agrobacterium-mediated transient expression

*A. tumefaciens* strains were grown on selective YEB plates and resuspended in infiltration buffer (10 mM MgCl_2_, 10 mM MES-KOH pH 5.6, 150 μM acetosyringone). Each strain was mixed with *A. tumefaciens* strain GV3101 pMP90 expressing the silencing suppressor 19K at a ratio of 1:1. The cultures were infiltrated into leaves of 4-5 week-old *N. benthamiana* plants using a needleless syringe. After 2 days, AvrRpm1 expression was induced by spraying infiltrated leaves with Dex as above and leaf material for protein extraction was harvested 8 h later.

### Plant protein extraction and detection by immunoblots

Protein extracts were prepared by grinding plant leaf material in liquid N_2_ to a fine powder followed by resuspension in extraction buffer [50 mM Tris-HCl, 150 mM NaCl, 10% (v/v) glycerol, 1 mM EDTA, 5 mM DTT, 1× protease inhibitor cocktail (Sigma-Aldrich #P9599), 0.2% NP-40, pH 7.4] at a ratio of 2 mL buffer per 1 g leaf material. Crude protein extracts were centrifuged at 20000 x *g* / 4 °C / 20 min and the supernatant was either used for SDS-PAGE or subjected to ADPr-enrichment with the Af1521 Macro domain. For immunoblots, proteins were separated by SDS-PAGE and electro-blotted onto PVDF membrane. The membrane was blocked with 5% non-fat dry milk in TBST for 1 h at RT. Proteins were detected using α-HA 3F10 antibody (Merck, 11867423001, 1:4000), α-Flag M2 antibody (Merck, F3165, 1:1000), α-pan-ADPr binding reagent (Merck, MABE1016, 1:4000), α-mono-ADPr AbD33204 (Bio-Rad, 1:2000), or α-ADPr E6F6A (Cell Signaling, #83732, 1:5000) in 2.5% non-fat dry milk in TBST over night at 4 °C. After three wash steps in TBST, membranes were incubated with HRP-coupled secondary antibodies (Merck, 1:20000). Proteins were detected using chemiluminescence substrates ECL Prime Western Blotting Detection Reagent (Cytiva) or SuperSignal™ West Femto Maximum Sensitivity Substrate (Thermo Fisher) and X-ray films (Fig. 1, Supplementary Fig. S5) or a Fusion Fx imager (Vilber) (Fig. 3, Supplementary Fig. S4).

### Enrichment of ADP-ribosylated proteins

Per affinity purification 15 μL Glutathione-Sepharose 4B resin (Merck) was washed in 50 mM Tris-HCl pH 7.5, 50 mM NaCl, 1 mM DTT and pelleted by centrifugation (500 x *g*, RT, 1 min). 1, 2.5 or 5 μg of GST-Macro (or G42E) were added in a total volume of 1 mL wash buffer and the tubes were rotated for 1 h at 4 °C. The resin was pelleted as above and washed twice with 0.8 mL plant extraction buffer (see plant protein extraction section above) supplemented with 10 μM MG132 (Merck). Cold plant extract was either added directly to the resin (Supplementary Dataset S1) or pre-cleared by filtration over 1-3 stacked 5 mL HiTrap desalting columns (Cytiva) followed by collection of the protein fraction based on absorbance at 280 nm (Supplementary Datasets S2-S8). The samples were rotated for 2 h at 4 °C and washed three times with 1 mL cold plant extraction buffer. The proteins were eluted from the resin by adding SDS sample buffer and incubation at 75 °C for 5 min.

### *In vitro* ADP-ribosylation assay

8 μg of His6-MKK5 [or His6-HaRxL106ΔC as control protein (Wirthmueller et al., 2015)] was diluted in reaction buffer (40 mM HEPES pH7.5, 5 mM MgCl_2_, 1 mM DTT, 30 μM NAD^+^, 60 μM ATP). Then either 4 μg His6-HopF2 or the corresponding volume of buffer (20 mM HEPES pH7.5, 150 mM NaCl) were added to a final volume of 50 μL. The reactions were incubated at 25 °C for 45 min. The samples were supplemented with 17 μL 4x SDS sample buffer and incubated at 75 °C for 3 min. ADP-ribosylation was detected by an immunoblot with α-ADPr (E6F6A) antibody as described above. As control, proteins were separated by SDS-PAGE and visualized by staining with Instant Blue (Abcam).

### Availability of data and constructs

The following plasmids have been submitted to Addgene (https://www.addgene.org/): pENTR4-MAMI #209186, pENTR4-PHOT1 #209187, pENTR4-PLDGAMMA3 #209188, pENTR4-AvrRpm1 #209185, pENTR4-HopU1 #228395, pOPINF-HopF2 #228397, pOPINF-MKK5 #228396. The pOPINF His6-GST-Af1521 Macro expression constructs are in part bound by a MTA with the University of Zurich and are, as all other materials, available upon request. The mass spectrometry proteomics data have been submitted to the ProteomeXchange Consortium via the PRIDE (Perez-Riverol et al., 2022) partner repository with the following identifiers: PXD045780 (Dataset S1); PXD045511, PXD045544, PXD045545 (Dataset S2); PXD045638, PXD045558, PXD045548 (Dataset S3); PXD045726 (Dataset S4); PXD045549 (Dataset S5); PXD056607 (Dataset S6), PXD056608 (Dataset S7), PXD045550 (Dataset S8).

## Acknowledgements

We thank the following colleagues for sharing published materials: Jeff Dangl (Dex-AvrRpm1 lines in *rpm1-3* background), Darrell Desveaux (Dex-HopF2 lines), Michael Hottiger (pGEX-6p-1 Af1521 K35E/Y145R Macro domain), Gitta Coaker (pTA7001/des/3XFlag). LW acknowledges core funding from the Leibniz Institute of Plant Biochemistry (IPB). We thank Jessica Erickson and Kee Hoon Sohn for helpful comments on the manuscript.

## Author contributions

SK, TC, and LW conceived and designed experiments. SK, DT, CP, and LW carried out experiments. SK, TC, SM, IM, and LW analyzed the data. LW wrote the manuscript with input from all co-authors. All authors reviewed and approved the submitted manuscript.

---

**Supplementary Figure S1.**
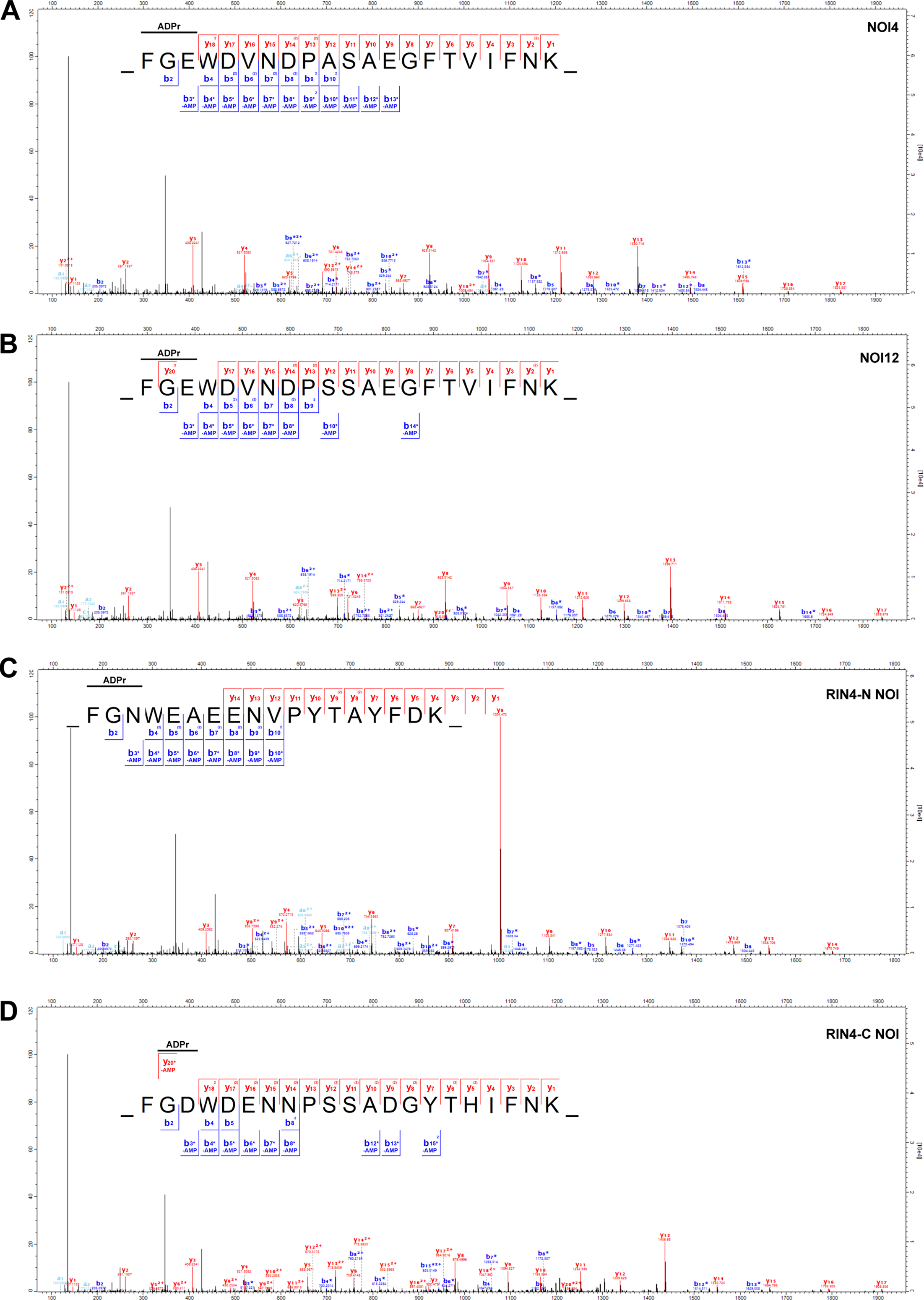
Analysis of MS^2^ spectra from the ADP-ribosylated NOI4, NOI12, and RIN4 peptides supports ADP-ribosylation on position 3 of the FGxW motif. Peak lists of the MS^2^ spectra shown in Fig. 2A-D were exported from Proteome Discoverer software and imported into the Expert system for computer-assisted annotation of MS/MS spectra (Neuhauser et al., 2012). Presence of the respective b3* ions with a neutral loss of AMP indicates ADP-ribosylation on E29 (NOI4, panel **A**), E13 (NOI12, panel **B**), N11 (RIN4 N-NOI, panel **C**), and D153 (RIN4 C-NOI, panel **D**) as F and G are unlikely to be ADP-ribosylated.

**Supplementary Figure S2.**
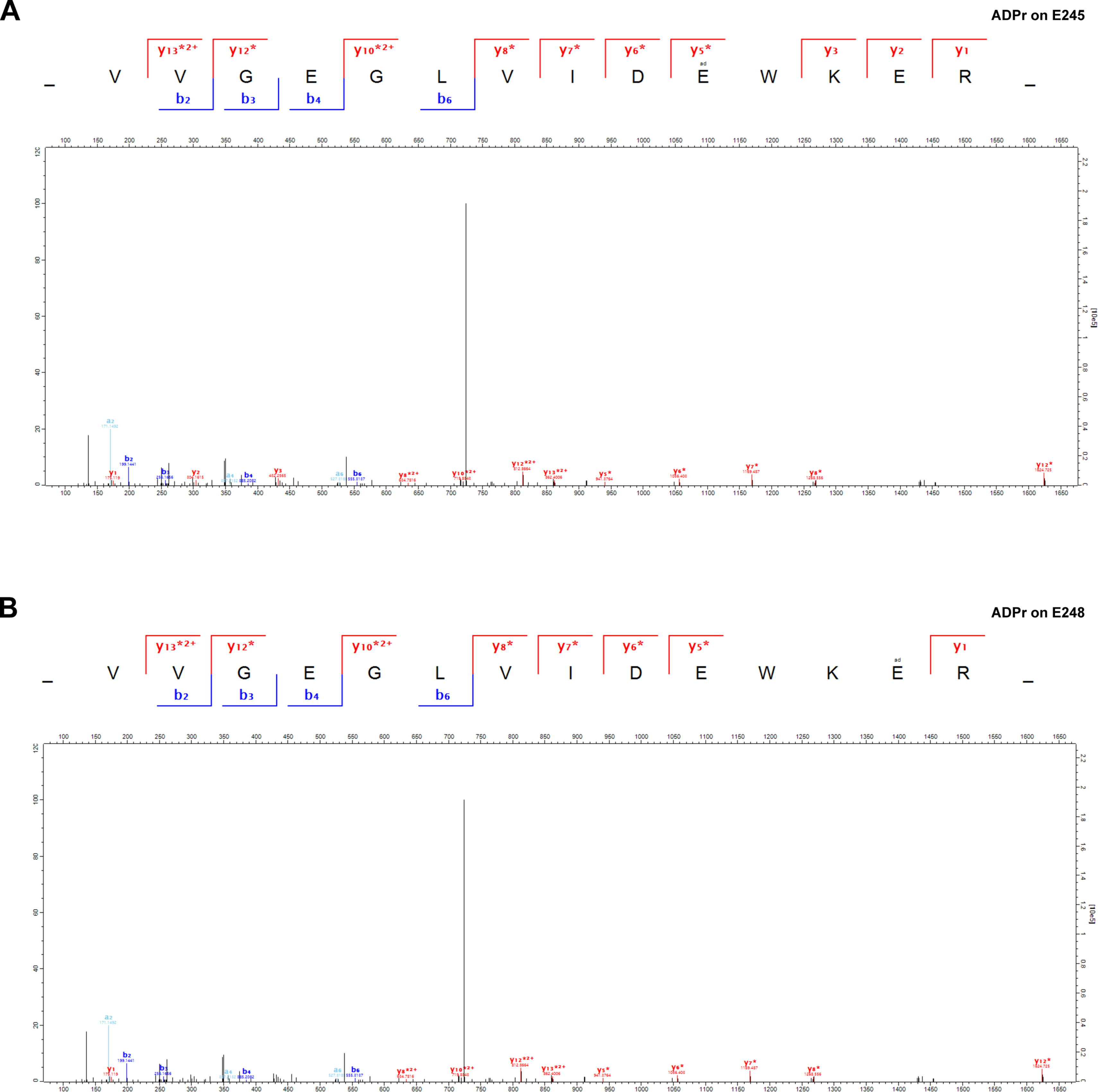
Analysis of the MS^2^ spectrum from the ADP-ribosylated MAMI peptide. The peak list of the MS^2^ spectrum shown in Fig. 3A was exported from Proteome Discoverer software and imported into the Expert system for computer-assisted annotation of MS/MS spectra (Neuhauser et al., 2012). **(A)** Match of the AMP neutral loss series assuming ADP-ribosylation on E245. The unmodified y2 and y3 ions are present. **(B)** Match of the AMP neutral loss series assuming ADP-ribosylation on E248. y2 and y3 ions with a neutral loss of AMP were not detected and ADPr on E248 does not allow annotation of additional peaks as compared to the model in (A).

**Supplementary Figure S3.**
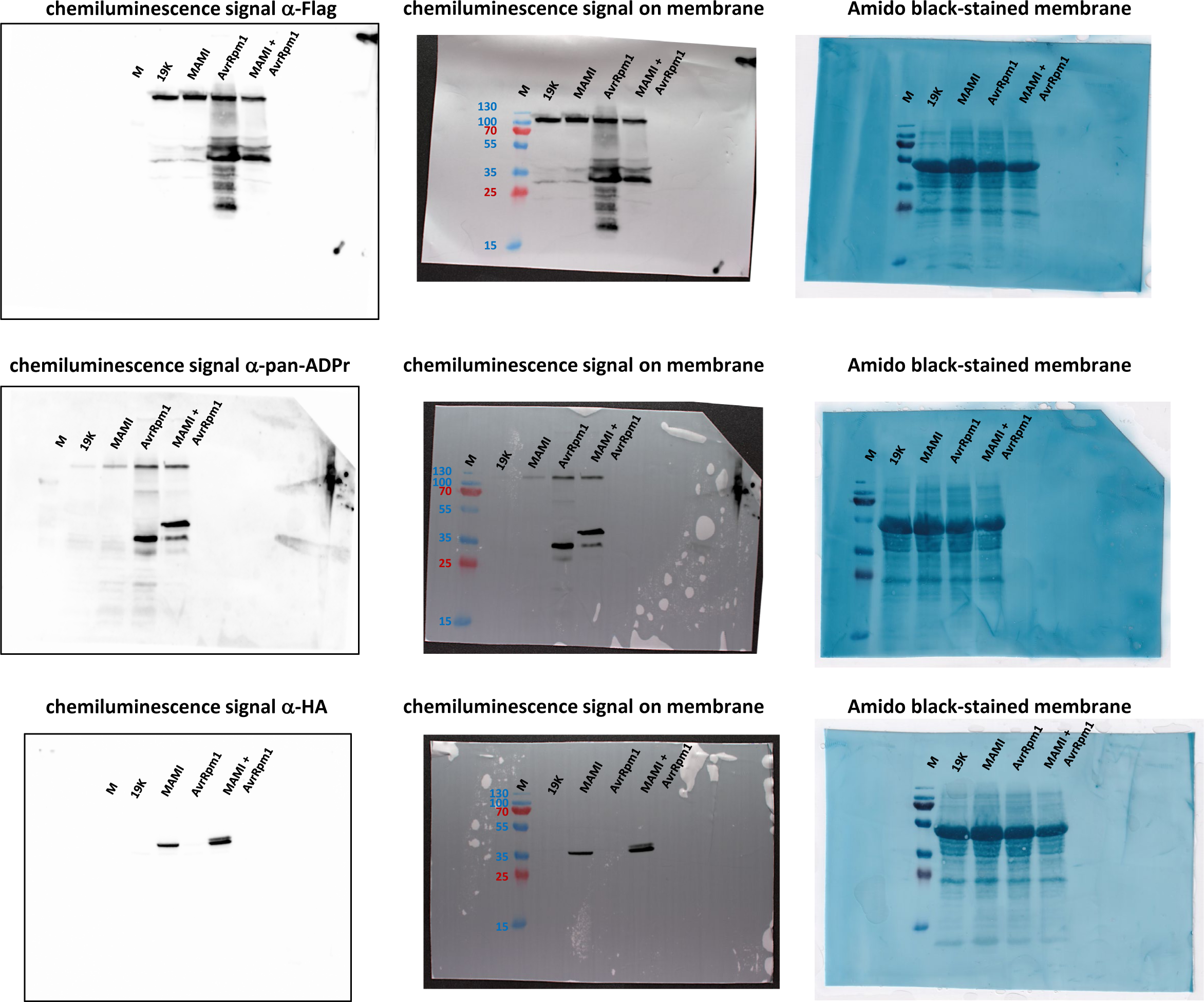
Images of the full membranes used for Figure 3B.

**Supplementary Figure S4.**
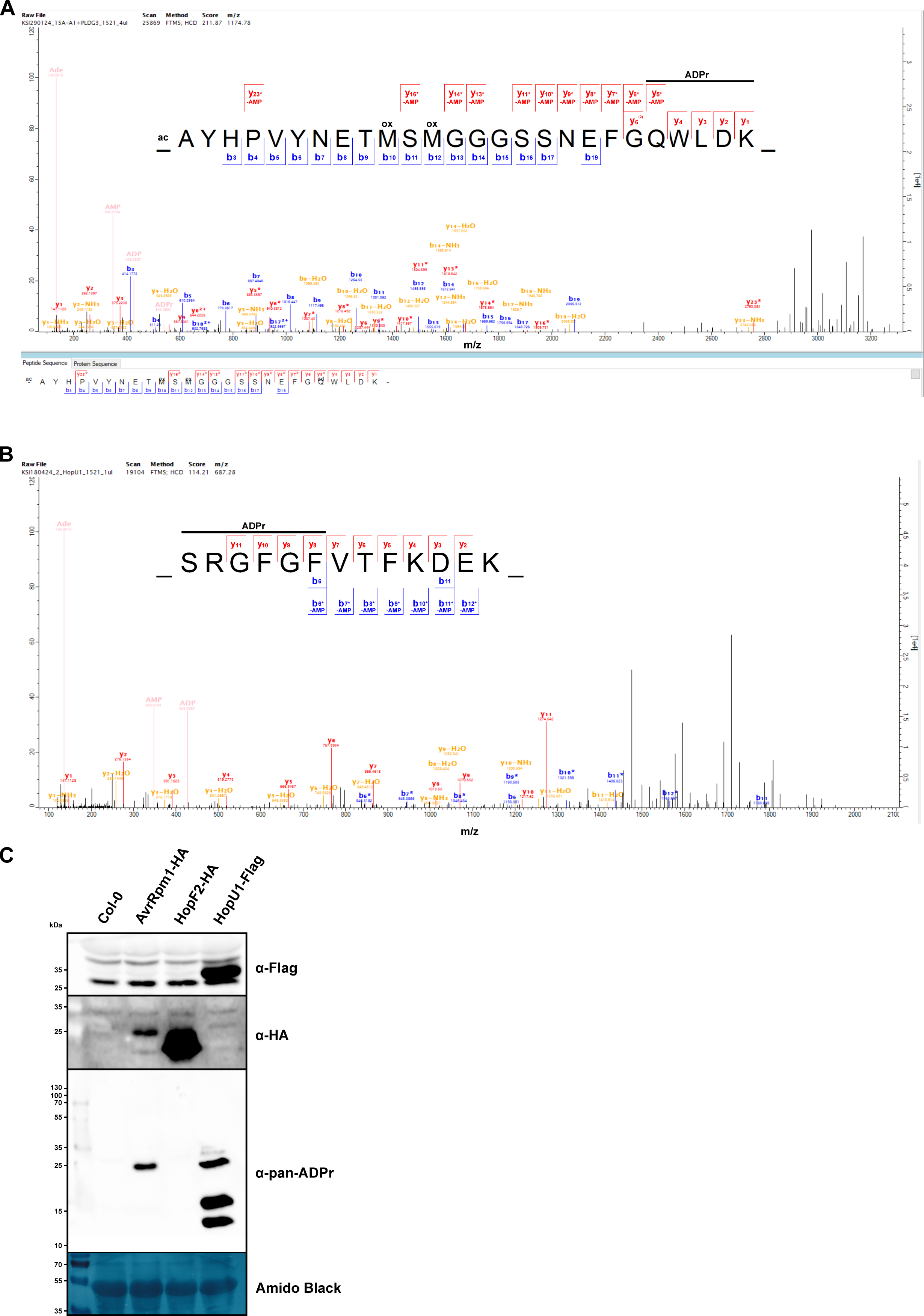
**(A)** HCD MS^2^ spectrum of the ADP-ribosylated peptide AYHPVYNETMSMGGGSSNEFGQWLDK (z = 3, mass = 3519.31 Da) from PLDGAMMA3-HA-StrepII co-expressed with AvrRpm1-Flag in *N. benthamiana*. **(B)** HCD MS^2^ spectrum of the ADP-ribosylated GRP7 peptide SRGFGFVTFKDEK (z = 3, mass = 2057.83 Da) identified from HopU1-expressing Arabidopsis plants. The ADPr marker ions are shown in beige. Unmodified fragment ions and fragment ions with a neutral loss of AMP (Δ347.06 Da) are indicated in the sequence logos. The horizontal line designates the possible sequence window for ADP-ribosylation. **(C)** Inducible expression of the T3E HopU1 results in ADP-ribosylation of proteins in Arabidopsis. Protein extracts from Dex-treated Col-0 wild type and the indicated transgenic lines were separated by SDS-PAGE, electroblotted onto PVDF membrane and probed with α-HA, a-Flag or α-pan-ADPr antibodies. The Amido Black-stained membrane is presented as loading control.

**Supplementary Figure S5.**
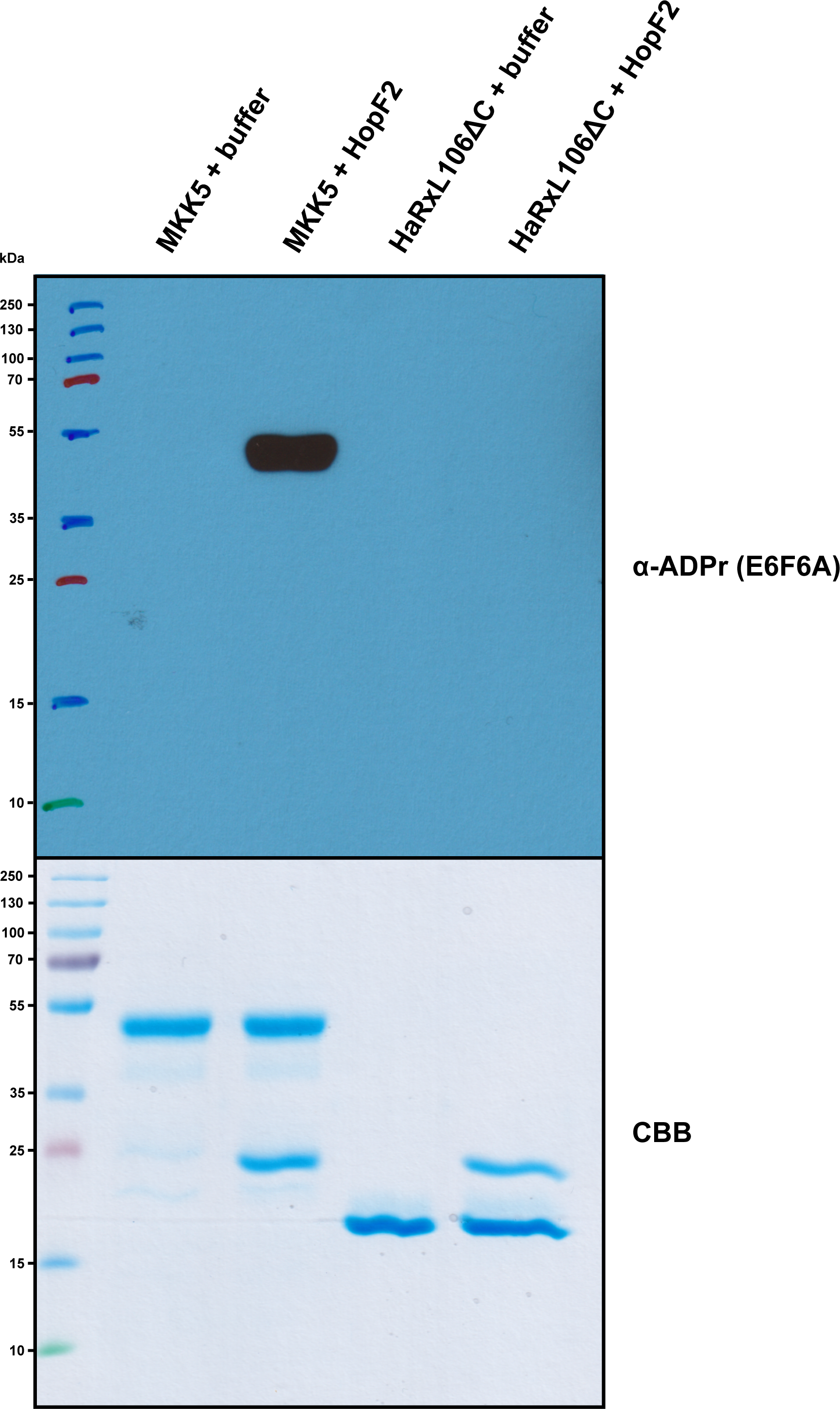
HopF2 can ADP-ribosylate MKK5 *in vitro*. His6-MKK5 or the control protein His6-HaRxL106ΔC were incubated with His6-HopF2 in ADP-ribosylation buffer at 25 °C for 45 min. In control reactions the equivalent volume of buffer was added instead of His6-HopF2. ADP-ribosylation was detected by immunoblotting with α-ADPr E6F6A antibody. As loading control, the same volume of the samples was analyzed by SDS-PAGE.

Supplementary Dataset S1. Proteins identified by untargeted proteomics in two Af1521 Macro pulldown experiments from Dex-treated Col-0, AvrRpm1-HA, and HopF2-HA plants.

Supplementary Dataset S2. ADP-ribosylated peptides identified in three Af1521 Macro pulldown experiments from the same AvrRpm1-HA sample. For ADPr-enrichment, either 1, 2.5 or 5 μg of the engineered Af1521 Macro domain was added to the protein extract. The data was analyzed with MaxQuant and the output file ‘ADP-ribosylationSites.txt’ and the information on the corresponding MS^2^ spectra from file ‘msms.txt’ were combined into one .xlsx file.

Supplementary Dataset S3. ADP-ribosylated peptides identified in three Af1521 Macro pulldown experiments from the same AvrRpm1-HA sample by the ‘triggering’ method. For ADPr-enrichment, either 1, 2.5 or 5 μg of the engineered Af1521 Macro domain was added to the protein extract. The data was analyzed with MaxQuant and the output file ‘ADP-ribosylationSites.txt’ and the information on the corresponding MS^2^ spectra from file ‘msms.txt’ were combined into one .xlsx file.

Supplementary Dataset S4. Comparison of proteins identified using the functional (Af1521) versus non-functional (G42E) Macro domains following ADPr-enrichment of proteins from Dex-treated AvrRpm1-HA plants.

Supplementary Dataset S5. *N. benthamiana* tissue expressing either AvrRpm1-Flag, StrepII-HA-MAMI, or co-expressing AvrRpm1-Flag and StrepII-HA-MAMI were subjected to ADPr-enrichment by the functional (Af1521) versus non-functional (G42E) Macro domains. The data was analyzed with MaxQuant and the output files ‘evidence.txt’, ‘ADP-ribosylationSites.txt’ and the information on the corresponding MS^2^ spectra from file ‘msms.txt’ were combined into one .xlsx file.

Supplementary Dataset S6. *N. benthamiana* tissue expressing either AvrRpm1-Flag, PLDGAMMA3-HA-StrepII, or co-expressing AvrRpm1-Flag and PLDGAMMA3-HA-StrepII were subjected to ADPr-enrichment by the functional (Af1521) versus non-functional (G42E) Macro domains. The data was analyzed with MaxQuant and the output files ‘evidence.txt’, ‘ADP-ribosylationSites.txt’ and the information on the corresponding MS^2^ spectra from file ‘msms.txt’ were combined into one .xlsx file.

Supplementary Dataset S7. ADP-ribosylated peptides identified in two Af1521 Macro pulldown experiments from HopU1-Flag vs. Col-0 samples by the ‘triggering’ method. Samples were subjected to ADPr-enrichment by the functional (Af1521) versus non-functional (G42E) Macro domains. The data was analyzed with MaxQuant and the output files ‘evidence.txt’, ‘ADP-ribosylationSites.txt’ and the information on the corresponding MS^2^ spectra from file ‘msms.txt’ were combined into one .xlsx file.

Supplementary Dataset S8. The data from Supplementary Dataset S3 were analyzed for phosphorylation on S and T and ADP-ribosylation using MaxQuant. The output files ‘Phospho (ST)Sites.txt’, ‘ADP-ribosylationSites.txt’ and the information on the corresponding MS^2^ spectra from file ‘msms.txt’ were combined into one .xlsx file.

Supplementary Dataset S9. Detailed description of proteomics methods used in this work.

Supplementary Methods.

## References

Aung, K., Kim, P., Li, Z., Joe, A., Kvitko, B., Alfano, J.R., and He, S.Y. (2020). Pathogenic Bacteria Target Plant Plasmodesmata to Colonize and Invade Surrounding Tissues. Plant Cell 32: 595–611.

Belkhadir, Y., Nimchuk, Z., Hubert, D.A., Mackey, D., and Dangl, J.L. (2004). Arabidopsis RIN4 negatively regulates disease resistance mediated by RPS2 and RPM1 downstream or independent of the NDR1 signal modulator and is not required for the virulence functions of bacterial type III effectors AvrRpt2 or AvrRpm1. Plant Cell 16: 2822–2835.

Berrow, N.S., Alderton, D., Sainsbury, S., Nettleship, J., Assenberg, R., Rahman, N., Stuart, D.I., and Owens, R.J. (2007). A versatile ligation-independent cloning method suitable for high-throughput expression screening applications. Nucleic Acids Res. 35: e45.

Bonfiglio, J.J., Leidecker, O., Dauben, H., Longarini, E.J., Colby, T., San Segundo-Acosta, P., Perez, K.A., and Matic, I. (2020). An HPF1/PARP1-Based Chemical Biology Strategy for Exploring ADP-Ribosylation. Cell 183: 1086–1102.

Chung, E.-H., da Cunha, L., Wu, A.-J., Gao, Z., Cherkis, K., Afzal, A.J., Mackey, D., and Dangl, J.L. (2011). Specific threonine phosphorylation of a host target by two unrelated type III effectors activates a host innate immune receptor in plants. Cell Host Microbe 9: 125–136.

Cunnac, S., Chakravarthy, S., Kvitko, B.H., Russell, A.B., Martin, G.B., and Collmer, A. (2011). Genetic disassembly and combinatorial reassembly identify a minimal functional repertoire of type III effectors in Pseudomonas syringae. Proc. Natl. Acad. Sci. U.S.A. 108: 2975–2980.

Dani, N., Stilla, A., Marchegiani, A., Tamburro, A., Till, S., Ladurner, A.G., Corda, D., and Di Girolamo, M. (2009). Combining affinity purification by ADP-ribose-binding macro domains with mass spectrometry to define the mammalian ADP-ribosyl proteome. Proc Natl Acad Sci U S A 106: 4243–4248.

Fu, Z.Q., Guo, M., Jeong, B., Tian, F., Elthon, T.E., Cerny, R.L., Staiger, D., and Alfano, J.R. (2007). A type III effector ADP-ribosylates RNA-binding proteins and quells plant immunity. Nature 447: 284–288.

Geiszler, D.J., Polasky, D.A., Yu, F., and Nesvizhskii, A.I. (2023). Detecting diagnostic features in MS/MS spectra of post-translationally modified peptides. Nat Commun 14: 4132.

He, Q., Naqvi, S., McLellan, H., Boevink, P.C., Champouret, N., Hein, I., and Birch, P.R.J. (2018). Plant pathogen effector utilizes host susceptibility factor NRL1 to degrade the immune regulator SWAP70. Proc Natl Acad Sci U S A 115: E7834–E7843.

Hendriks, I.A., Larsen, S.C., and Nielsen, M.L. (2019). An Advanced Strategy for Comprehensive Profiling of ADP-ribosylation Sites Using Mass Spectrometry-based Proteomics. Mol Cell Proteomics 18: 1010–1026.

Jeong, B., Lin, Y., Joe, A., Guo, M., Korneli, C., Yang, H., Wang, P., Yu, M., Cerny, R.L., Staiger, D., Alfano, J.R., and Xu, Y. (2011). Structure function analysis of an ADP-ribosyltransferase type III effector and its RNA-binding target in plant immunity. J. Biol. Chem. 286: 43272–43281.

Karras, G.I., Kustatscher, G., Buhecha, H.R., Allen, M.D., Pugieux, C., Sait, F., Bycroft, M., and Ladurner, A.G. (2005). The macro domain is an ADP-ribose binding module. EMBO J 24: 1911–1920.

Li, W., Chiang, Y.-H., and Coaker, G. (2013). The HopQ1 Effector’s Nucleoside Hydrolase-Like Domain Is Required for Bacterial Virulence in Arabidopsis and Tomato, but Not Host Recognition in Tobacco. PLoS One 8: e59684.

Liu, J., Elmore, J.M., Lin, Z.-J.D., and Coaker, G. (2011). A receptor-like cytoplasmic kinase phosphorylates the host target RIN4, leading to the activation of a plant innate immune receptor. Cell Host Microbe 9: 137–146.

Mackey, D., Holt, B.F., Wiig, A., and Dangl, J.L. (2002). RIN4 interacts with Pseudomonas syringae type III effector molecules and is required for RPM1-mediated resistance in Arabidopsis. Cell 108: 743–754.

Martello, R., Leutert, M., Jungmichel, S., Bilan, V., Larsen, S.C., Young, C., Hottiger, M.O., and Nielsen, M.L. (2016). Proteome-wide identification of the endogenous ADP-ribosylome of mammalian cells and tissue. Nat Commun 7: 12917.

Martin, R., Qi, T., Zhang, H., Liu, F., King, M., Toth, C., Nogales, E., and Staskawicz, B.J. (2020). Structure of the activated ROQ1 resistosome directly recognizing the pathogen effector XopQ. Science 370: eabd9993.

Naqvi, S., He, Q., Trusch, F., Qiu, H., Pham, J., Sun, Q., Christie, J.M., Gilroy, E.M., and Birch, P.R.J. (2022). Blue-light receptor phototropin 1 suppresses immunity to promote Phytophthora infestans infection. New Phytol 233: 2282–2293.

Neuhauser, N., Michalski, A., Cox, J., and Mann, M. (2012). Expert system for computer-assisted annotation of MS/MS spectra. Mol Cell Proteomics 11: 1500–1509.

Nowak, K., Rosenthal, F., Karlberg, T., Bütepage, M., Thorsell, A.-G., Dreier, B., Grossmann, J., Sobek, J., Imhof, R., Lüscher, B., Schüler, H., Plückthun, A., Leslie Pedrioli, D.M., and Hottiger, M.O. (2020). Engineering Af1521 improves ADP-ribose binding and identification of ADP-ribosylated proteins. Nat Commun 11: 5199.

Perez-Riverol, Y., Bai, J., Bandla, C., García-Seisdedos, D., Hewapathirana, S., Kamatchinathan, S., Kundu, D.J., Prakash, A., Frericks-Zipper, A., Eisenacher, M., Walzer, M., Wang, S., Brazma, A., and Vizcaíno, J.A. (2022). The PRIDE database resources in 2022: a hub for mass spectrometry-based proteomics evidences. Nucleic Acids Res 50: D543–D552.

Rack, J.G.M., Palazzo, L., and Ahel, I. (2020). (ADP-ribosyl)hydrolases: structure, function, and biology. Genes Dev 34: 263–284.

Redditt, T.J., Chung, E.-H., Karimi, H.Z., Rodibaugh, N., Zhang, Y., Trinidad, J.C., Kim, J.H., Zhou, Q., Shen, M., Dangl, J.L., Mackey, D., and Innes, R.W. (2019). AvrRpm1 Functions as an ADP-Ribosyl Transferase to Modify NOI Domain-Containing Proteins, Including Arabidopsis and Soybean RPM1-Interacting Protein4. Plant Cell 31: 2664–2681.

Rosenthal, F., Nanni, P., Barkow-Oesterreicher, S., and Hottiger, M.O. (2015). Optimization of LTQ-Orbitrap Mass Spectrometer Parameters for the Identification of ADP-Ribosylation Sites. J Proteome Res 14: 4072–4079.

Saur, I.M.L., Panstruga, R., and Schulze-Lefert, P. (2021). NOD-like receptor-mediated plant immunity: from structure to cell death. Nat Rev Immunol 21: 305–318.

Schöning, J.C., Streitner, C., Page, D.R., Hennig, S., Uchida, K., Wolf, E., Furuya, M., and Staiger, D. (2007). Auto-regulation of the circadian slave oscillator component AtGRP7 and regulation of its targets is impaired by a single RNA recognition motif point mutation. Plant J. 52: 1119–1130.

Seong, K. and Krasileva, K.V. (2021). Computational Structural Genomics Unravels Common Folds and Novel Families in the Secretome of Fungal Phytopathogen Magnaporthe oryzae. Mol Plant Microbe Interact 34: 1267–1280.

Suskiewicz, M.J., Prokhorova, E., Rack, J.G.M., and Ahel, I. (2023). ADP-ribosylation from molecular mechanisms to therapeutic implications. Cell 186: 4475–4495.

Tyanova, S., Temu, T., and Cox, J. (2016). The MaxQuant computational platform for mass spectrometry-based shotgun proteomics. Nat Protoc 11: 2301–2319.

Wang, J., Hu, M., Wang, J., Qi, J., Han, Z., Wang, G., Qi, Y., Wang, H.-W., Zhou, J.-M., and Chai, J. (2019). Reconstitution and structure of a plant NLR resistosome conferring immunity. Science 364: eaav5870.

Wang, Y., Li, J., Hou, S., Wang, X., Li, Y., Ren, D., Chen, S., Tang, X., and Zhou, J.-M. (2010). A Pseudomonas syringae ADP-ribosyltransferase inhibits Arabidopsis mitogen-activated protein kinase kinases. Plant Cell 22: 2033–2044.

Wang, Y., Pruitt, R.N., Nürnberger, T., and Wang, Y. (2022). Evasion of plant immunity by microbial pathogens. Nat Rev Microbiol 20: 449–464.

Willems, P., Horne, A., Van Parys, T., Goormachtig, S., De Smet, I., Botzki, A., Van Breusegem, F., and Gevaert, K. (2019). The Plant PTM Viewer, a central resource for exploring plant protein modifications. Plant J. 99: 752–762.

Wilton, M., Subramaniam, R., Elmore, J., Felsensteiner, C., Coaker, G., and Desveaux, D. (2010). The type III effector HopF2Pto targets Arabidopsis RIN4 protein to promote Pseudomonas syringae virulence. Proc. Natl. Acad. Sci. U.S.A. 107: 2349–2354.

Wirthmueller, L., Roth, C., Fabro, G., Caillaud, M.-C., Rallapalli, G., Asai, S., Sklenar, J., Jones, A.M.E., Wiermer, M., Jones, J.D.G., and Banfield, M.J. (2015). Probing formation of cargo/importin-α transport complexes in plant cells using a pathogen effector. Plant J. 81: 40–52.

Wu, W., Zou, H., Zheng, H., Chen, X., Luo, X., Fan, X., Zhuo, T., and Miao, W. (2024). Ralstonia solanacearum type III effector RipAF1 mediates plant resistance signaling by ADP-ribosylation of host FBN1. Hortic Res 11: uhae162.

Xin, X.-F., Nomura, K., Aung, K., Velásquez, A.C., Yao, J., Boutrot, F., Chang, J.H., Zipfel, C., and He, S.Y. (2016). Bacteria establish an aqueous living space in plants crucial for virulence. Nature 539: 524–529.

Yoon, M., Middleditch, M.J., and Rikkerink, E.H.A. (2022). A conserved glutamate residue in RPM1-INTERACTING PROTEIN4 is ADP-ribosylated by the Pseudomonas effector AvrRpm2 to activate RPM1-mediated plant resistance. Plant Cell 34: 4950–4972.

Yotsui, I., Matsui, H., Miyauchi, S., Iwakawa, H., Melkonian, K., Schlüter, T., Michavila, S., Kanazawa, T., Nomura, Y., Stolze, S.C., Jeon, H.-W., Yan, Y., Harzen, A., Sugano, S.S., Shirakawa, M., Nishihama, R., Ichihashi, Y., Ibanez, S.G., Shirasu, K., Ueda, T., Kohchi, T., and Nakagami, H. (2023). LysM-mediated signaling in Marchantia polymorpha highlights the conservation of pattern-triggered immunity in land plants. Curr Biol 33: 3732–3746.e8.

Zhou, J., Wu, S., Chen, X., Liu, C., Sheen, J., Shan, L., and He, P. (2014). The Pseudomonas syringae effector HopF2 suppresses Arabidopsis immunity by targeting BAK1. Plant J 77: 235–245.

