## Supplementary Methods for "Untargeted proteomics identifies plant substrates of the bacterial-derived ADP-ribosyltransferase AvrRpm1"

Protein expression and purification

His6-GST-Macro variants, His6-HopF2, and His6-MKK5 were expressed in *E. coli* Shuffle cells (NEB). Cultures were grown in LB medium at a temperature of 37 °C to an OD_600_ of 1.0 – 1.2. The cultures were cooled to 18 °C before expression was induced by the addition of 0.5 mM IPTG for 16 h. Cells were pelleted by centrifugation (5000 x *g* / 4 °C / 12 min) and the pellets were resuspended in buffer A [50 mM Tris-HCl, 0.3 M NaCl, 20 mM imidazole, 5% (v/v) glycerol, 50 mM glycine, pH 8.0] supplemented with 0.1% polyethylenimine and 1x cOmplete™ EDTA-free protease inhibitor cocktail (Merck). Cells lysis was induced by addition of Lysozyme (1 mg/mL final concentration / 25 °C / 15 min) followed by sonication on ice (Branson 150D Sonifier, 2x 10 min, level 3-4). Insoluble proteins and cell debris were removed by centrifugation (30000 x *g* / 4 °C / 30 min) and the supernatant was loaded onto a 5 mL HisTrap HP IMAC column (Cytiva). The column was washed with buffer A until the A_280_ reached 25 mAU and proteins were eluted using buffer B [50 mM Tris-HCl, 0.3 M NaCl, 0.5 M imidazole, 5% (v/v) glycerol, 50 mM glycine, pH 8.0]. The elution from the IMAC column was injected onto a size exclusion chromatography column [Superdex 75 26/60 PG column (Cytiva) pre-equilibrated with 20 mM HEPES-NaOH, 150 mM NaC, pH 7.5]. Proteins eluting from the column were concentrated by ultrafiltration on Vivaspin 20 columns (Sartorius) with a 5 kDa molecular weight cut-off. For His6-MKK5 we selected the peak corresponding to the monomeric form. For the His6-GST-Macro variants the tag was cleaved using 3C protease. The protein was run through a 5 mL HisTrap HP IMAC column in buffer A to remove the His6-tag and residual un-cleaved fusion protein, followed by injection onto the Superdex 75 26/60 PG column and elution and concentration as above. Aliquots were snap-frozen in liquid N_2_ and stored at -70 °C.

Sample preparation for LC-MS/MS

Protein samples were separated by 12% SDS-PAGE until the running front had migrated 1 cm into the separating gel. The gels were stained with Coomassie and the protein area was cut from the gel. Proteins were in-gel digested with trypsin and desalted as described by (Majovsky et al., 2014).

Acquisition of raw data sets by higher energy collisional dissociation (HCD) fragmentation

Peptides were analyzed with 5 different MS methods on a Fusion Lumos Tribrid mass spectrometer (Thermo Fisher Scientific), summarized in Supplementary Dataset S9. Dried peptides were dissolved in 5% acetonitrile, 0.1% trifluoric acid and injected into an EASY-nLC 1200 liquid chromatography system (Thermo Fisher Scientific). Peptides were separated using liquid chromatography C18 reverse phase chemistry employing a 120 min gradient increasing from 1 to 24 % acetonitrile in 0.1% FA (MS method 1) or from 1% to 36% acetonitrile in 0.1% FA (MS methods 2-5), and a flow rate of 250 nL/min. Eluted peptides were electrosprayed on-line into the Fusion Lumos Tribrid with a spray voltage of 2.0 kV and capillary temperature of 305°C. Peptides (MS^1^) were detected in the Orbitrap with the following settings: resolving power: 120,000; scan range m/z 300–1500; S-Lens RF 30%; AGC target Standard, max injection time (IT) on Auto mode, microscans: 1. Dynamic exclusion duration was set for 60 s (MS method 1) or 30 s (MS methods 2-5). MS/MS peptide sequencing was performed using a Top15 DDA scan strategy (MS method 1), or peptides were selected for a 1 s time window in between master scans in a data-dependent mode (MS methods 2-5) with HCD at 30% NCE, to be detected in the Orbitrap with a resolution of 30000. For differences in AGC target and maximum injection time mode between different methods see Supplementary Dataset S9. Different targeted mass exclusion lists containing the most prominent peptide m/z ratios for the GST-Af1521 Macro domain and, where appropriate, GSTs from *A. thaliana* were used for MS methods 2, 3 and 5 (see Supplementary Dataset S9). For Methods 3-5, an additional MS^2^ scan with an isolation window of 1.3 was triggered by the presence of the diagnostic peak of adenine at 136.062 Da in a precursor’s fragmentation spectrum. Orbitrap detection occurred at a resolution of 60000 with a defined first mass of 120 m/z. The normalized AGC Target of the triggered scan was set to 1000% and the maximum injection time was 1800 ms.

Identification of proteins and peptides

For Supplementary Datasets S1 and S4, peptides and proteins were identified using the Mascot software v2.7.0 (Matrix Science) linked to Proteome Discoverer v2.4 (Thermo Fisher Scientific). The enzyme specificity was set to trypsin and two missed cleavages were tolerated. Carbamidomethylation of cysteine (C) was set as fixed modification, and oxidation of methionine (M) as well as ADP-ribosylation of aspartic acid (D) or glutamic acid (E) or phosphorylation of serine (S) and threonine (T) as variable modifications. A precursor ion mass error of 10 ppm and a fragment ion mass error of 0.02 Da were tolerated in searches of the TAIR10 database amended with sequences of GST-Af1521 and common contaminants. A decoy database search was performed to determine the peptide spectral match (PSM) and peptide identification false discovery rates (FDR). Peptides with a score surpassing the false discovery rate threshold of 0.01 (q-value<0.01) were considered positive identifications. The phosphoRS module was used to specifically map ADP-ribosylation or phosphorylation to amino acid residues within the primary structure of peptides.

For Supplementary Datasets S2, S3, S5, S6, S7, and S8 peptides and proteins were identified using MaxQuant software v2.1.3.0 (Tyanova et al., 2016). For Supplementary Datasets S2, S3, S7, and S8 the data were searched against the *A. thaliana* Araport11 proteome database downloaded from <https://www.arabidopsis.org/download/> on 03.01.2022 and the sequences of the GST-Af1521 Macro domains. For Supplementary Datasets S5 and S6 peptides and proteins were identified by searches against the *N. benthamiana* NbDE proteome database (Kourelis et al., 2019) downloaded on 22.12.2022. In all searches a FDR of 0.01 was applied for PSM, protein, and site decoy fraction. The minimal peptide length was 7 amino acids, the enzyme specificity was set to Trypsin/P and two missed cleavages were tolerated. Carbamidomethylation of cysteine (C) was set as fixed modification. Variable modifications were oxidation of methionine (M), acetylated protein N-termini and ADP-ribosylation of aspartic acid (D), glutamic acid (E), asparagine (N), lysine (K), arginine (R), serine (S), cysteine (C), and histidine (H). For PLDGAMMA3 expressed in *N. benthamiana* (Dataset S6) ADP-ribosylation of glutamine (Q) was added in addition. Neutral losses of Adenine (135.0545 Da), AMP (347.0631 Da), Adenosine H_2_O (249.0862 Da), ADP (427.0294 Da), and ADPr (541.0611 Da) were included for data analysis. Adenine^+^ (136.0618 Da), AMP^+^ (348.0704 Da), and ADPr^+^ (542.0684 Da) were used as diagnostic peaks for ADP-ribosylation. For Supplementary Dataset S6 phosphorylation of serine (S) and threonine (T) was included as additional variable modification. For analysis of ADP-ribosylated peptides in MaxQuant we i) applied an Andromeda score of >80 and ii) manually inspected MS^2^ spectra for ADPr marker ions and neutral losses characteristic of ADP-ribosylation.

**References**

**Kourelis, J., Kaschani, F., Grosse-Holz, F.M., Homma, F., Kaiser, M., and van der Hoorn, R.A.L.** (2019). A homology-guided, genome-based proteome for improved proteomics in the alloploid Nicotiana benthamiana. BMC Genomics **20**: 722.

**Majovsky, P., Naumann, C., Lee, C.-W., Lassowskat, I., Trujillo, M., Dissmeyer, N., and Hoehenwarter, W.** (2014). Targeted proteomics analysis of protein degradation in plant signaling on an LTQ-Orbitrap mass spectrometer. J Proteome Res **13**: 4246–4258.

**Tyanova, S., Temu, T., and Cox, J.** (2016). The MaxQuant computational platform for mass spectrometry-based shotgun proteomics. Nat Protoc **11**: 2301–2319.
